# Orthogonal Representations of Reward Magnitude, Certainty, and Volatility in the Macaque Orbitofrontal Cortex

**DOI:** 10.1101/365080

**Authors:** Tianming Yang, Elisabeth A. Murray

## Abstract

Categorical knowledge about the probabilistic and volatile nature of resource availability can improve foraging strategies, yet we have little understanding of how the brain represents such knowledge. Neurons in the orbitofrontal cortex (OFC) of macaques encode several decision variables (e.g., reward magnitude, probability) that could influence choice behavior. Here we investigated whether OFC neurons also represent two aspects of reward predictability: certainty and volatility. Rhesus monkeys performed a visual stimulus-reward association task in which a set of simple shapes preceded the delivery of reward, and they learned the nature of each shape’s reward association along two dimensions. One involved the certainty of a reward outcome; rewards can be either deterministic (and therefore certain) or probabilistic (uncertain). A second dimension reflected the volatility of an outcome; reward schedules can be either stable over time or volatile. During stimulus presentation, the activity of OFC neurons reflected both categorical certainty and categorical volatility, in addition to reward magnitude. These three characteristics were represented orthogonally by three distinct neural populations of similar size. These findings point to a more general role for OFC in processing reward information than one restricted to encoding parametric valuations such as reward magnitude and probability.

---

Foraging decisions take into account several aspects of reward, including reward quantity, quality, availability and costs, among other factors, yet decision-making would benefit from knowledge of more abstract aspects of reward, such as certainty and volatility. Knowledge of certainty and volatility helps to interpret uncertain outcomes. For example, a slot machine player may repeat his bet many times even if he is currently losing, knowing that one cannot be certain of the outcome of a single play, and the odds of winning are fixed. But a poker player often adjusts his strategy because he knows his opponents’ playing strategy may change over the course of play.

In the present study, we considered two categorical reward attributes that affect how we adapt our behavior based on reward feedback. The first attribute is certainty, defined as whether the stimulus-reward association is probabilistic. One may not be certain of the reward outcome if it is probabilistic. The second is volatility, which indicates a change over time. According to our definitions, a stimulus-reward association can be both certain and volatile, meaning it is stable and non-probabilistic during a period of time; in such cases a change in the association can be detected with confidence.

In a static environment, neurons in the orbitofrontal cortex (OFC) have been shown to encode many aspects of reward, including magnitude (Wallis and Miller 2003), probability (Kennerley and Wallis 2009), risk (O’Neill and Schultz 2015), abstract rules (Wallis et al. 2001), and strategies (Tsujimoto et al. 2011). More importantly, the OFC has been identified as essential for adaptive behavior when the environment is volatile. Lesions of the OFC lead to severe deficits when stimulus-outcome contingencies need to be updated (Jones and Mishkin 1972; Dias et al. 1996; Izquierdo et al. 2004; Walton et al. 2010). Although recent work with selective excitotoxic lesions of the OFC in macaques (Rudebeck et al. 2013) has overturned the idea that the OFC is necessary for learning reversals of stimulus-reward contingencies—at least in deterministic settings—recordings from single neurons have shown that OFC neurons change their activity to visual stimuli when the type of their associated outcome changes (Thorpe et al. 1983; Rolls et al. 1996; Morrison et al. 2011) and during learning (Wallis and Miller 2003; Rudebeck et al. 2017). Taken together, these results suggest the OFC plays an important role in tracking stimulus-reward associations.

In the current study, we build on previous studies of the OFC and ask the question whether categorical information of certainty and volatility of reward is encoded in the OFC in addition to reward magnitude. Because certainty and volatility are tightly related to value, it is possible that these three attributes are combined and represented together in single OFC neurons. Alternatively, however, OFC neurons might encode some task-relevant parameters separately, as seems to be the case for risk and value (O’Neill and Schultz 2015) and information and value (Blanchard et al. 2015).

To test these ideas, we trained two monkeys on a task in which the stimulus-outcome associations differ in certainty and volatility. A set of well-learned simple geometric shapes were provided as cues to indicate the nature of the association. The monkeys had to employ their knowledge of certainty and volatility aspects of the stimulus-outcome associations to predict rewards. Consistent with earlier work, we found that many neurons in OFC encoded the reward magnitude associated with each stimulus. In addition, we found neurons that indicated whether the cue association was certain or uncertain, and stable or volatile. We found no support for the idea that reward magnitude, certainty, and volatility are combined at the level of single OFC neurons. Instead, these three attributes were represented orthogonally by different groups of neurons.

## Materials and Methods

### Subjects

Two male rhesus monkeys (*Macaca mulatta*), naïve at the start of the experiment, were used in the study. They weighed 7.8 and 8.5 kg at the beginning of the experiment. All experimental procedures conformed to the Guide for the Care and Use of Laboratory Animals and were approved by the National Institute of Mental Health Animal Care and Use Committee.

### Materials

Monkeys were seated in a primate chair facing a 15-inch video monitor. Stimuli subtending 1 degree of visual angle were displayed on the monitor screen at central fixation. Rewards consisted of 1 to 3 drops of juice (0.01~0.2 ml per drop), with drop size controlled by a pressurized reward delivery system (Precision Engineering Co., MD, USA). Eye position was monitored with an infrared occulometer (Arrington Research).

### Behavioral Task

We trained two monkeys to perform a visual cue-reward association task (Figure 1A). Each trial began with the appearance of a fixation point in the center of the screen. When the monkey gazed at the fixation point and simultaneously held its hand on a centrally located button (left button in Figure 1A), a single geometric shape was presented on the monitor for a variable period of time ranging between 500 and 1000 ms. The cue predicted either 1 drop or 3 drops of juice. Each drop of juice was 0.1~0.2 ml. When the cue was extinguished, the monkey had to move its hand to another button on the right and hold it there. A touch to the right button led to the appearance of a green square in the center of the screen. If the monkey expected to receive one drop of juice, then it needed to hold the right button for 1-2 seconds. If the monkey expected 3 drops of juice, it needed to hold for 3-4 seconds. If the monkey held the right button for the correct duration, the green square disappeared and the juice was delivered immediately. If the monkey released the button too soon or too late, it would receive the same number of drops of juice, but each drop was only one-tenth of that received for a correct response, specifically, 0.01~0.02 ml. In addition, if the monkey released the right button too fast, the green square would turn orange and the monkey was required to wait until the correct hold time was fulfilled, plus a 2-second penalty, before reward was delivered. For example, if the correct response was to hold for 3 seconds, and the monkey released the button at 2.5 seconds, then it had to wait 0.5 plus 2 seconds, or a total of 2.5 seconds, before reward delivery. The intertrial interval was 1 second.

**Figure 1.**
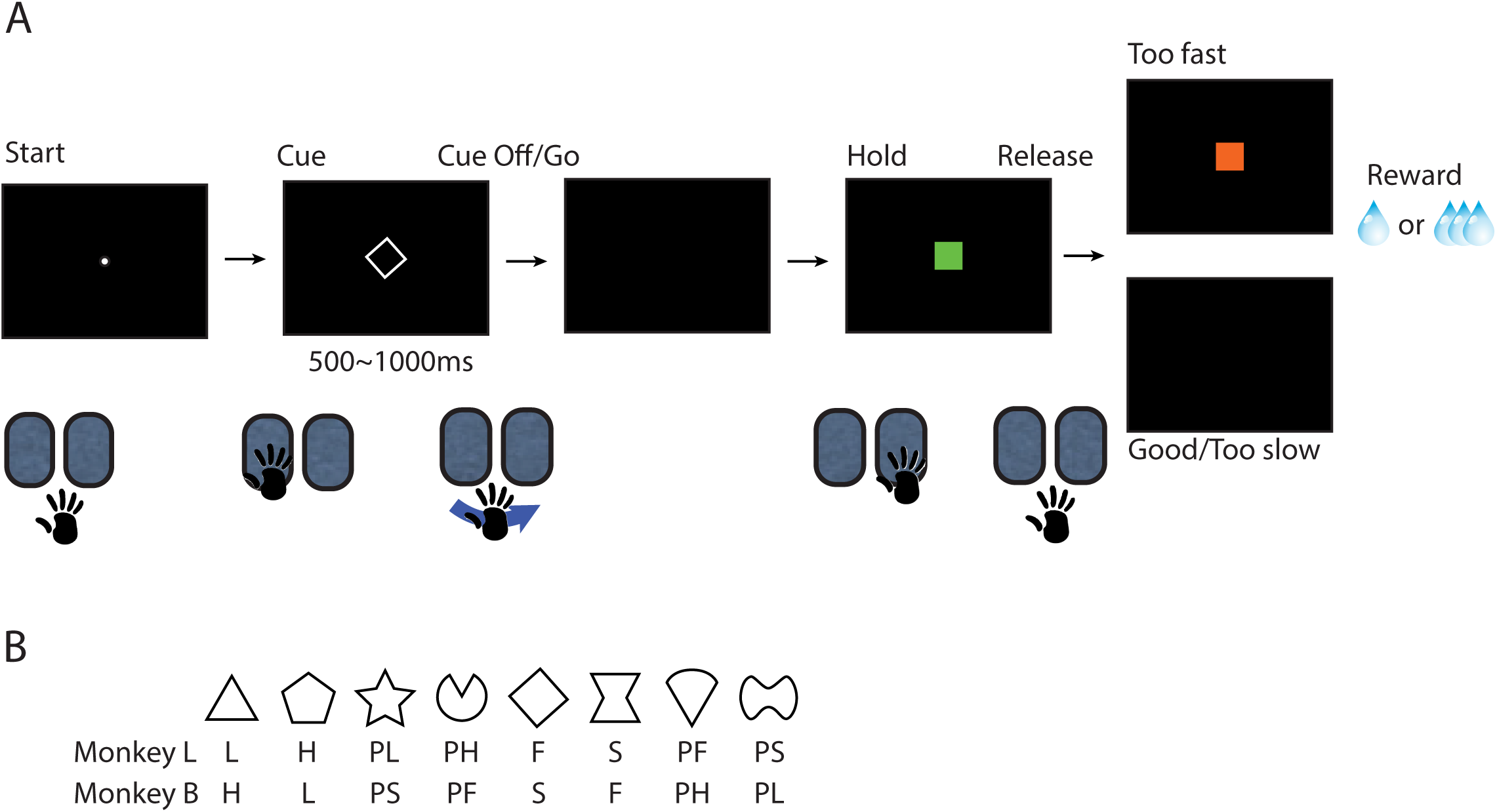
**A**. Trial sequence. A central dot on the monitor screen indicated the initiation of a new trial. Once the monkey touched the left button, a single visual cue out of a set of 8 appeared on the screen. When the cue was turned off, the monkey had to remove its hand from the left button and place it on the right button. A green square appeared when the monkey’s hand touched the right button. The monkey had to hold its hand on the right button for 1~2 seconds if it expected to receive one drop of juice, and 3~4 seconds if it expected to receive 3 drops of juice. An orange square appeared if the monkey released the button too fast, and the orange square stayed on the screen for an extra 3 seconds as a penalty. If the monkey released too late, the green square would be extinguished as the time expired. The same number of drops of juice would be delivered regardless of the monkey’s response. However, if the monkey’s holding time was too short or too long, it only received drops of juice that were 1/10 of the size that it received for correct responses. **B**. The cue-reward assignments. L: certain 1 drop; H: certain 3 drops; PL: fixed probabilistic (uncertain) with 1 drop being more likely; PH: fixed probabilistic (uncertain) with 3 drops being more likely; S: certain but slow-changing; F: certain but fast-changing; PS: probabilistic (uncertain) slow-changing; PF: probabilistic (uncertain) fast-changing.

The visual cues informed the monkey of the number of drops of upcoming juice. A single set of 8 visual cues was used (Figure 1B), including 2 certain cues that predicted fixed amounts of 1 drop (low, L) and 3 drops (high, H) of juice, 2 uncertain and probabilistic cues that predicted 3 drops of juice at fixed probabilities 2/3 (PH) and 1/3 (PL), and 2 volatile but certain cues that predicted juice that would change its number of drops between 1 drop and 3 drops of juice at the probability of 0.02 and on average every 50 trials (fast, F) and at the probability of 0.005 and on average every 200 trials (slow, S). Finally, 2 volatile and uncertain cues signaled that reward outcomes would change from 3 drops on 1/3 of trials to 3 drops on 2/3 of trials at the probability of 0.02 and on average every 50 trials (PF) and at the probability of 0.005 and on average every 200 trials (PS). The cue-state transitions were randomly determined using the specified probabilities at each trial. When a cue switched its reward association, there was a 5-trial window in which its cue-reward contingency was kept the same. The cue-reward correspondence was assigned differently between the two monkeys.

We started training the animals with the two stable and certain cues, then added the stable and uncertain cues, the volatile and certain cues, and, finally, the volatile and uncertain cues, in that order. At each stage, we made sure the animals’ performance was over 70% correct before introducing the new cues. In the final stage of training as well as during the recording sessions, the cue presented on each trial was selected randomly from the set of 8.

### Surgery

After initial training, a titanium headpost was implanted on the cranium of each monkey using standard surgical procedures. The monkeys were then retrained on the task. When task performance was satisfactory, a second operation was performed to implant an acrylic recording chamber over the prefrontal region. A craniotomy was created inside the chamber.

All operations were performed under aseptic conditions. Monkeys were immobilized with ketamine hydrochloride (10 mg/kg, i.m.). Glycopyrrolate was administered to reduce secretions (0.01 mg/kg, i.m.). During surgery, anesthesia was induced and maintained with isoflurane gas (1-3%, to effect). Isotonic fluids were given via an intravenous catheter. Body temperature, heart rate, blood pressure, and expired CO_2_ were monitored throughout the surgical procedures. Cefazolin antibiotic (15 mg/kg, i.m.) was given one day before surgery and for one week after to prevent infection. In addition, monkeys received the analgesic ketoprofen (10-15 mg, i.m.) at the end of surgery and for two additional days. This was followed by 100 mg of ibuprofen for the next 5 days.

### MRI

After the recording chamber was implanted, and after the first few recordings were performed, we acquired structural magnetic resonance imaging (MRI) scans in a 3T scanner to verify the recording locations. Monkeys were sedated with a mixture of ketamine (15-20 mg/kg, i.m.) and medetomidine (20 µg/kg, i.m.), supplemented as needed. Glycopyrrolate was administered to reduce secretions (0.01 mg/kg, i.m.) and ketoprofen (10-15 mg, i.m.) was given as an analgesic. Monkeys were placed in a MR-compatible stereotaxic frame for the duration of the scan.

### Electrophysiology

We recorded single unit activity with vertically movable electrodes (FHC or Alpha Omega, 0.5–1.5 MΩ at 1 KHz) using conventional techniques. Briefly, microelectrodes were driven by an eight-channel micromanipulator (NAN Instruments) attached to the recording chamber. We used at most four electrodes at the same time. Recordings locations were on the ventral surface of the frontal lobe between the lateral and medial orbital sulci, roughly corresponding to Walker’s (1940) areas 11 and 13 (Figure 2). Spike waveforms from putative single neurons were isolated online and recorded with a multi-channel acquisition processor (Plexon). Spike sorting was carried out offline (Plexon off-line sorter). Other than the quality of isolation, there were no selection criteria for neurons. We confirmed recording locations in monkey B by placing marking lesions and performing histological analyses at the end of the experiment (Figure 2).

**Figure 2.**
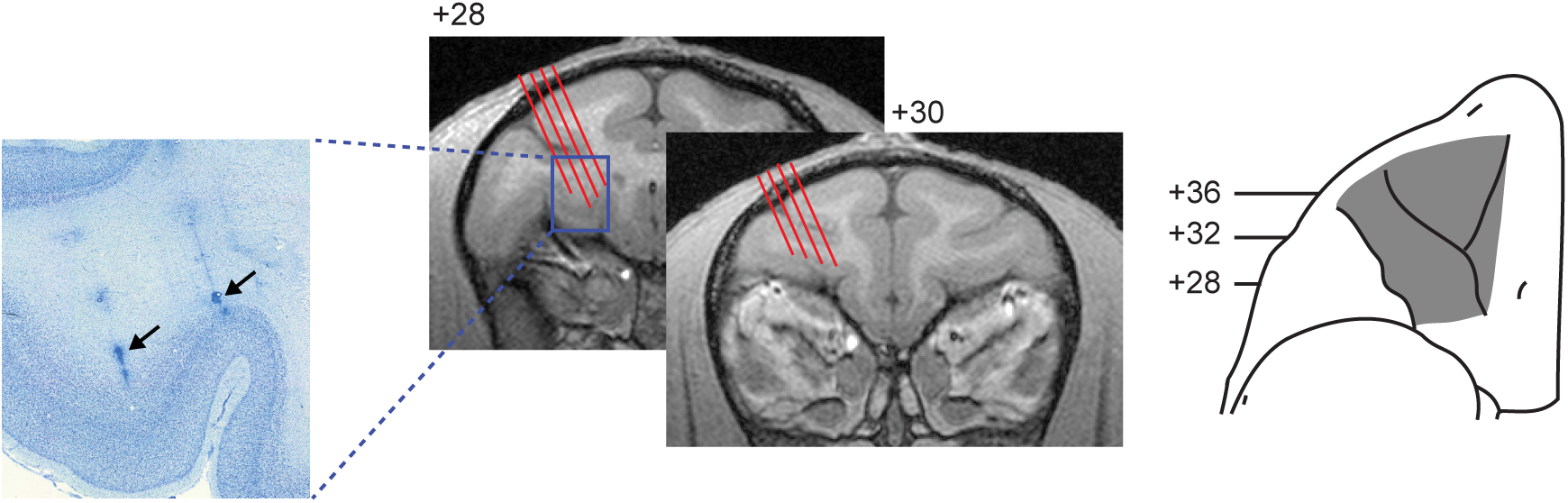
Recording locations in monkey B. Electrode tracks (red) and locations in the OFC are shown schematically on T1-weighted MRI coronal sections. Numerals indicate distance in millimeters from the interaural plane. Black arrows on photomicrograph (left) indicate marking lesions that were used to reconfirm the recording locations; shading on the ventral view of the frontal lobe (right) shows the approximate region from which neurons were recorded.

### Behavioral Data Analysis

All behavioral analyses were based on the monkeys’ behavior during the recording sessions. The trials are the same ones that we used in analyzing the activity from single neurons.

#### Accuracy

We defined the monkeys’ responses as correct when they held the button for 3 seconds for certain cues that led to 3 drops of juice or uncertain cues that were more likely to yield 3 drops of juice. Similarly, the monkeys’ responses were considered correct when they held the button for 1 second for certain cues that led to 1 drop of juice or uncertain cues that were more likely to yield 1 drop of juice.

#### Reward History Analysis

We estimated the influence of past rewards delivered in the trials of a particular cue on monkeys’ holding time of this cue in the current trial by regressing the holding time against past rewards:

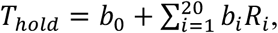

where *R_i_* indicates the reward of the *i-*th trials with the same cue back in the history, with 0 indicating a reward of 1 drop of juice and 1 indicating a reward of 3 drops of juice. The history extends up to 20 trials back without additional transitions. Values of *b_1_*… *b*_20_ indicated the influence of past rewards on monkeys’ current prediction of reward associated with the cue and were plotted in Figure 3C.

**Figure 3.**
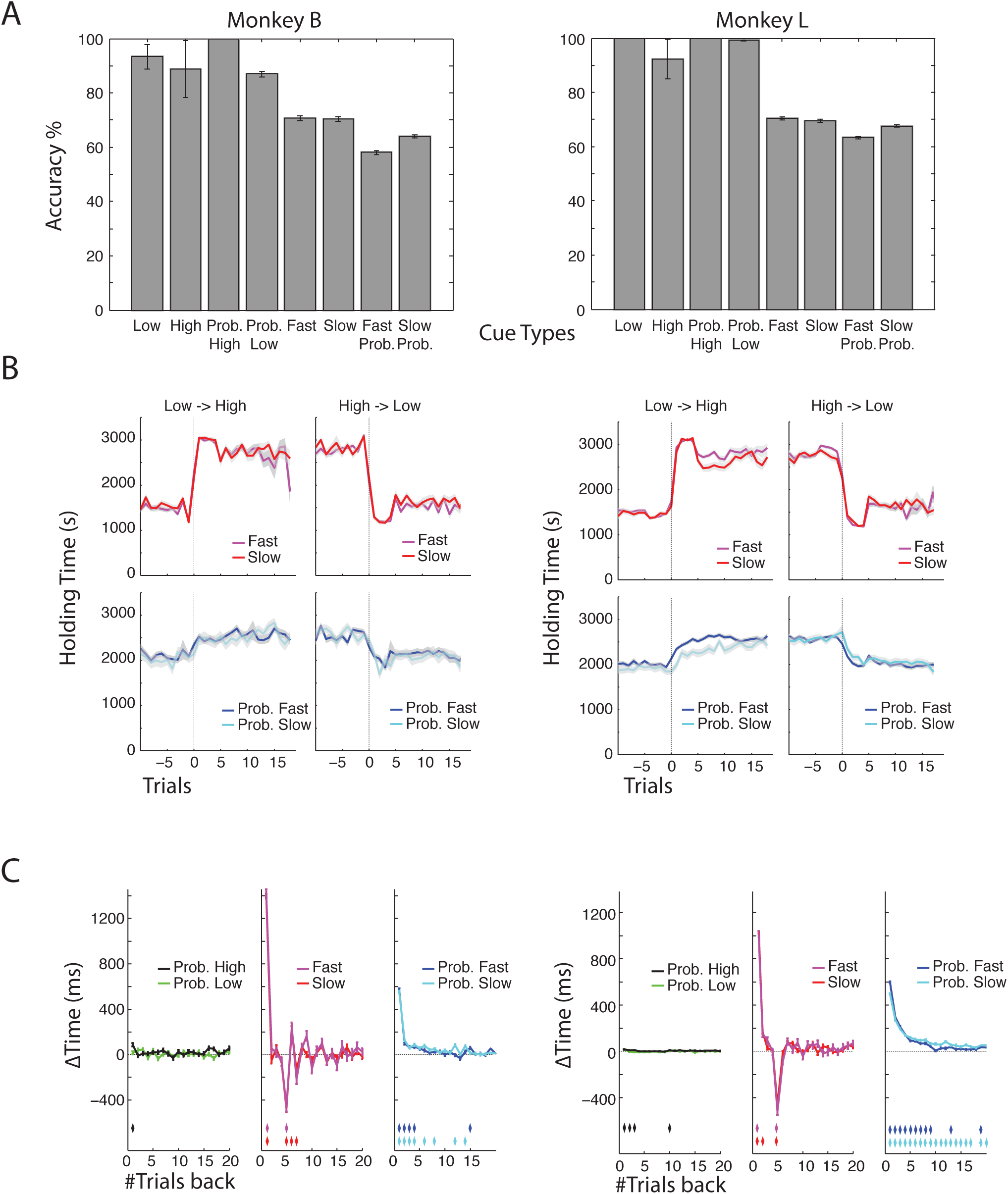
The choice behavior of monkeys B and L. **A**. The monkeys’ performance for each cue type averaged across sessions. The error bars indicate Bernoulli distribution standard error of the mean. **B**. The holding time for each cue type at the time of transition. Curves show the mean (line) and standard errors (shaded area). The top row: certain cues (F and S), the bottom row: uncertain cues (PF and PS). **C**. The time added to the current trial’s holding time by 3-drops-of-juice trials in the past. The error bars indicate the 95% confidence intervals. Trials that differ significantly from 0 after Bonferroni correction are indicated by the diamonds with corresponding colors in the bottom of each plot.

#### Neural Data Analysis

We determined each neuron’s tuning properties using a regression:

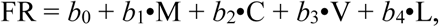

where FR is the measured firing rate of the neuron in a trial, M indicates whether the monkeys predicted the cue’s reward outcome was 1 or 3 drops of juice, C indicates whether the stimulus-reward association was certain or not, V indicates whether the stimulus-reward association was volatile or not, and L indicates whether the monkey received 1 drop or 3 drops of juice in the last trial. The cue period firing rate was the neuron’s average response within a 500-ms period after the cue onset. The pre-release period firing rate was calculated using a 500-ms time window before the release of the right button, using the correct trials only. The significance was determined at p<0.05.

Time-course analyses were performed by using a 5-ms sliding window (Figures 4–6). The population response was done by combining all trials from all neurons in subpopulations selected based on the regression results (Figures 5 and 6). The significance of response difference (Figure 6) was determined with a t-test for each time window at p<0.05 with Bonferroni correction. The latency of the population encoding of each tuning property was defined as the start of a period of at least 10 continuous 5-ms time bins (50 ms in total) with a significant difference after the stimulus onset. To verify that our results were not artifacts generated from certain neurons with high firing rates or with a lot of trials, we performed analyses using a variety of normalization methods and obtained qualitatively very similar results. The tuning indices for each neuron (Figure 5) were calculated as following:

**Figure 4.**
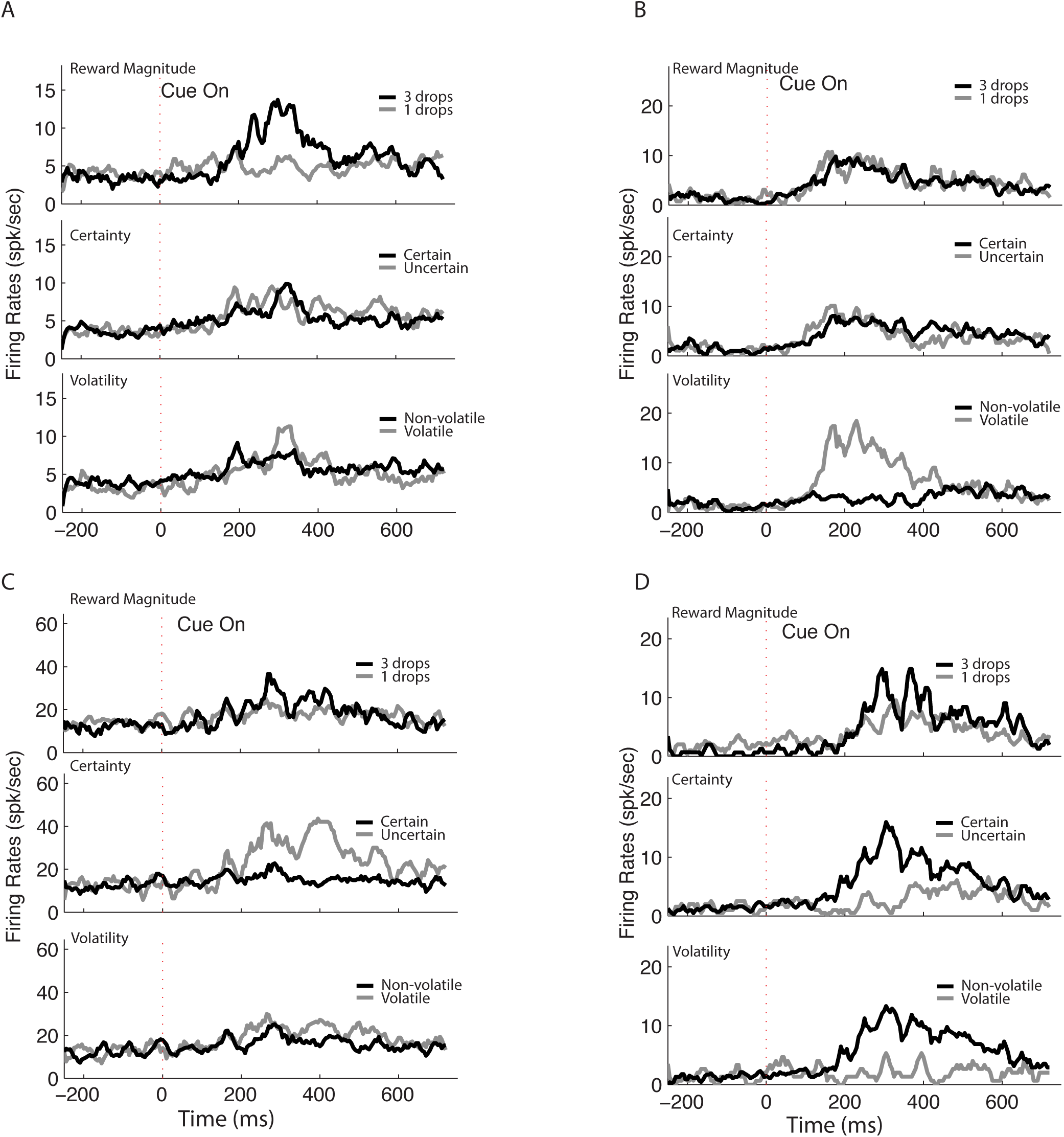
The cue-period responses for some example neurons: **A**. A neuron tuned to reward magnitude. **B**. A neuron tuned to volatility. **C**. A neuron tuned to certainty. **D**. A neuron with mixed tuning. The classification is based on the activity of each neuron during the 500-ms period after cue onset.

**Figure 5.**
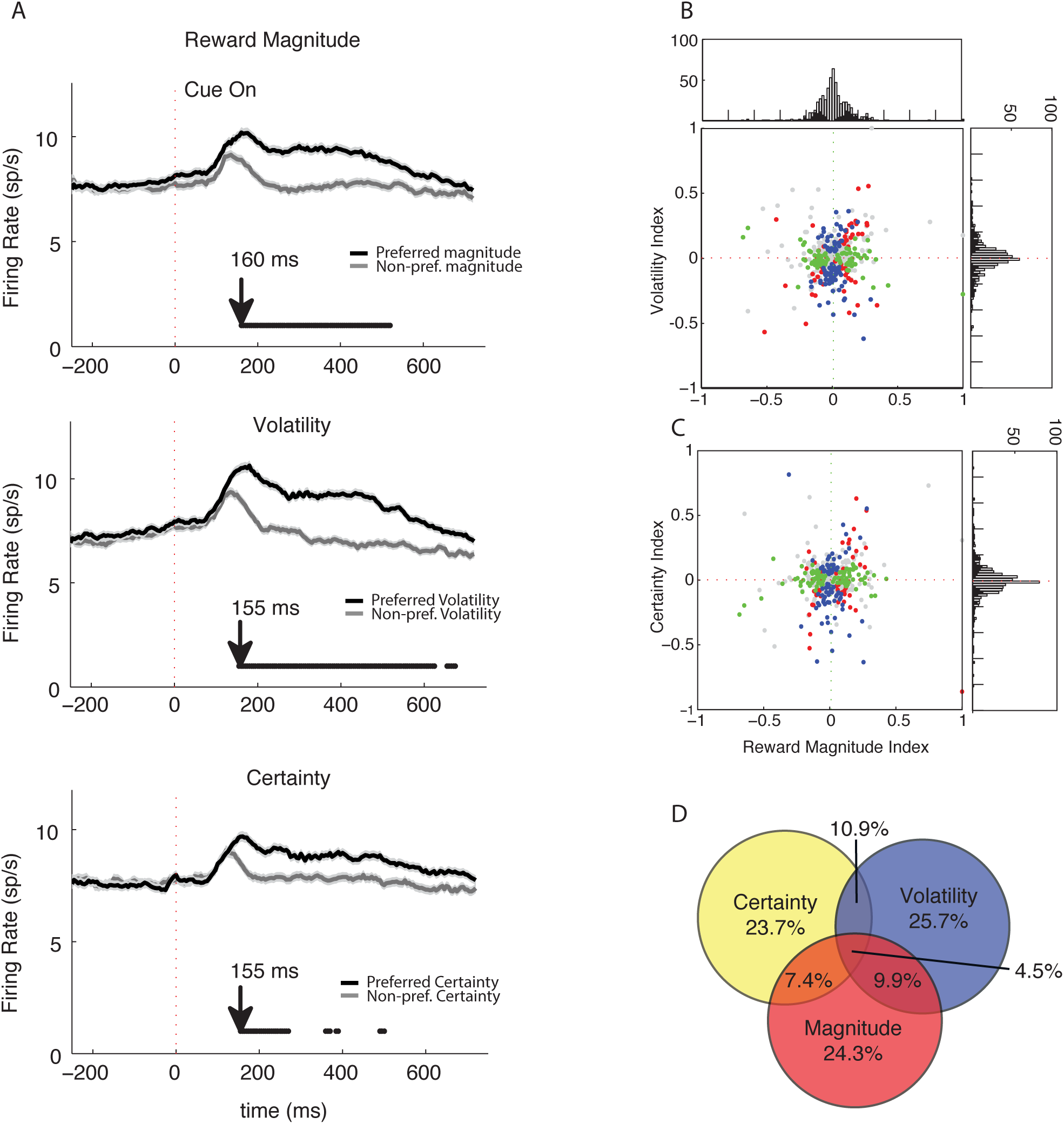
**A**. OFC neurons’ population responses to visual cues sorted on the cues’ reward magnitude, certainty and volatility. That is, the classification used here is based on the results of the regression and classification as reported in Table 1. Arrows indicate the latency at which a tuning property is significant for the population. **B**. The volatility indices during the cue period of each neuron plotted against the reward magnitude indices. Each dot is a neuron. Blue indicates neurons with significant volatility tuning. Green indicates neurons with significant reward magnitude tuning. Red indicates neurons tuned to both. Grey indicates neurons with no significant tuning. The distribution of reward magnitude indices is plotted in the horizontal histogram on the top, with the black bars indicating significant tuning. The volatility index distribution was similarly plotted in the vertical histogram on the right. **C**. The certainty indices plotted against the reward magnitude indices in a similar format as in panel B. Blue indicates neurons with significant certainty tuning. Green indicates neurons with significant reward magnitude tuning. Red indicates neurons tuned to both. Grey indicates neurons with no significant tuning. The certainty index distribution is plotted in the vertical histogram on the right. **D**. The Venn diagram of the neurons that are tuned to magnitude, certainty and volatility.

**Figure 6.**
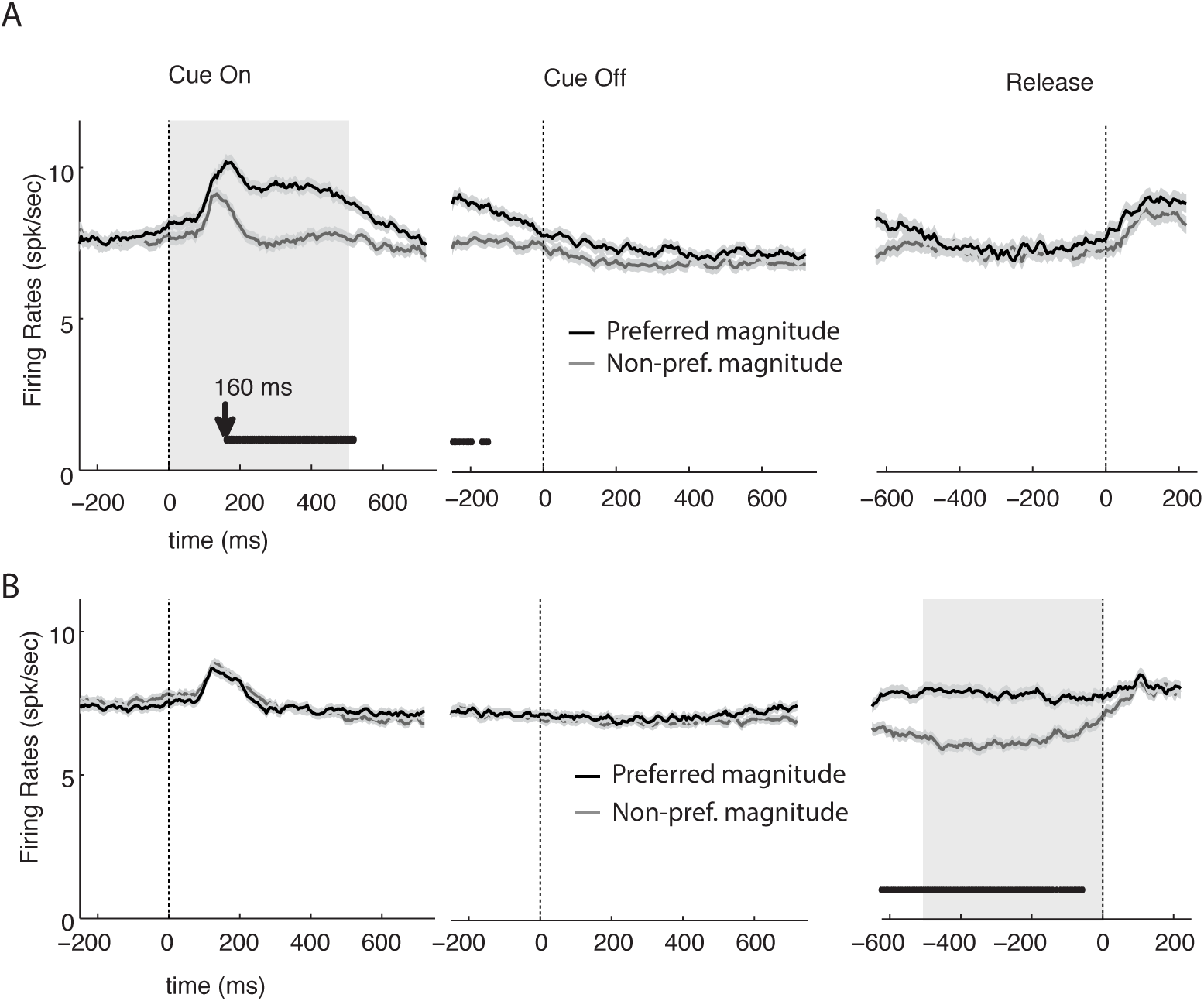
Inconsistent population encoding of reward magnitude between the cue period and pre-release period. **A.** The population responses of the OFC neurons that are reward magnitude tuned during the cue period are aligned to the cue onset, cue offset and the animals’ response, signaled by release of the right button. The dark lines near the X-axis indicate the time during which the two curves are significantly different. The gray box indicates the window used to evaluate the tuning. Arrow indicates the latency at which magnitude tuning is significant for the population. **B.** The population responses of the OFC neurons that are reward magnitude tuned during the 500-ms period before the right button release are aligned to the cue onset, cue offset and the animals’ response. The gray box indicates the window used to evaluate the tuning.

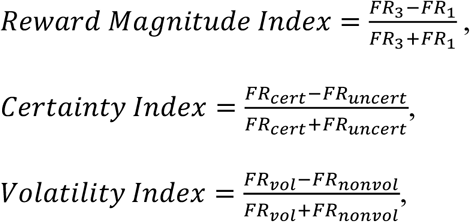

where *FR*_3_ and *FR*_1_ are the neuron’s firing rates within the period under consideration when the monkey predicted that the cue would lead to 3 and 1 drop(s) of juice, respectively; *FR*_cert_ and *FR*_uncert_ are when the cue was certain and uncertain, respectively; and *FR*_vol_ and *FR*_nonvol_ are when the cue was volatile and nonvolatile respectively.

## Results

### Behavior

We trained two monkeys to perform a visual stimulus-reward association task (Figure 1). Briefly, on each trial a single visual stimulus from a fixed set of 8 was presented on a monitor screen. When a ‘go signal’ was given, the monkey was required to press a button for either 1 second or 3 seconds. To earn the maximum amount of reward, the monkey needed to apply a hold period that indicated the expected reward magnitude: 1 second for 1 drop and 3 seconds for 3 drops. Because four of the cues were volatile and varied their reward associations during the session, the monkeys had to track those associations and adjust their responses accordingly.

The optimal choice strategy for each cue is different. The stable certain cues demand monkeys to have a fixed response throughout the experiment. For the stable and uncertain cues, the best strategy is to select a hold time that reflects the more likely outcome always, i.e., holding the button for 3 seconds if 3 drops of juice is the more likely outcome, and holding for 1 second if 1 drop is the more likely outcome. For the volatile but certain cues, the best strategy is a win-stay, lose-shift strategy using only the outcome of the last trial. The most complicated case is the cues that are both volatile and uncertain. Monkeys cannot predict the current state with just the last trial’s outcome. They need to integrate over a period time to deduce the current state and make their estimate of reward size accordingly.

After extensive training, both monkeys were able to perform the task (Figure 3A). For the stable cues, they were able to hold the button for an appropriate length of time on over 90% of the trials, regardless whether they were certain or not. For the volatile but certain cues, their performance was still very good; monkeys applied the appropriate hold time on roughly 70% of trials. Monkeys’ performance with the volatile and uncertain cues was, understandably, lower; each monkey applied the correct hold times on close to 60% of trials, which was still significantly above chance levels (p<0.05).

To find out how the monkeys reacted to the changes of stimulus-reward associations, we analyzed their behavior around the time of transitions. For the volatile and certain cues, the monkeys adjusted their behavior almost immediately after the transition. In contrast, the strategy adjustment happened much more gradually for the uncertain ones (Figure 3B). These results were confirmed by calculating the amount of influence of past trials on the monkeys’ hold times (Figure 3C). For the volatile and certain cues, only the last trial’s outcome had significant effects on the current trial hold time, and the effect was large. For the volatile and uncertain cues, a long history of recent trial outcomes affected the current trial hold time. The history influence was strongest for the last trial and dropped off gradually back in the history. Neither monkey’s responses for the uncertain but stable cues were affected by the reward history.

In summary, the monkeys learned to track the reward information in a manner consistent with what we would expect from the optimal strategy, suggesting they had learned the categorical certainty and volatility information of the cues and used the knowledge to track and estimate their reward association.

### OFC electrophysiology

Neurons were recorded in lateral OFC areas 11 and 13 (Walker 1940; Carmichael and Price 1994). We recorded from 370 neurons from monkey L and 279 neurons from monkey B. For the purpose of analysis, neurons from both regions and both monkeys were considered together. Numbers from each monkey are provided in Table 1. They show the same trends as the combined data.

**Table 1.** Numbers of neurons that are modulated by specific reward attributes in the cue period and pre-release period. For each period, the first row shows the total, and the second and third rows show the numbers from monkeys L and B respectively. Considered attributes include magnitude (Mag.), certainty (Cert.), volatility (Vol.), all combinations of the three, and reward in the previous trial (Prev Rew). Neurons that are not modulated by any of the above attributes are also shown (None).

|  | Mon<br>keys | Mag. | Cert. | Vol. | Mag. &<br>Cert. | Mag. &<br>Vol. | Cert. &<br>Vol. | All<br>three | Prev<br>Rew | None | Total |
| --- | --- | --- | --- | --- | --- | --- | --- | --- | --- | --- | --- |
| Cue<br>period | Both | 158 | 154 | 167 | 48 | 64 | 71 | 29 | 148 | 253 | 649 |
|  | L | 76 | 81 | 83 | 21 | 23 | 33 | 11 | 84 | 150 | 370 |
|  | B | 82 | 86 | 71 | 27 | 41 | 38 | 18 | 64 | 103 | 279 |
| Pre-release<br>period | Both | 244 | 101 | 81 | 42 | 43 | 22 | 13 | 77 | 279 | 649 |
|  | L | 138 | 49 | 69 | 27 | 28 | 13 | 9 | 46 | 151 | 370 |
|  | B | 106 | 32 | 32 | 15 | 15 | 9 | 4 | 31 | 128 | 279 |

We focused our analysis on the three variables signaled by the visual cues: reward magnitude, volatility, and certainty. Reward magnitude is a binary variable indicating the association in a given trial between the visual cue and the reward, whether it is 1 drop or 3 drops of juice. For many cues, reward magnitude dynamically varies within a session. The holding time of the monkey indicates the monkey’s predicted reward magnitude. Both volatility and certainty are also binary variables indicating whether a cue is volatile and whether it is certain. Half of our 8 shapes were volatile (F, S, PF, and PS, see Figure 3B), and half of them were probabilistic and thus uncertain (PL, PH, PF, and PS), with 2 of them being both (PF and PS), and 2 of them being neither (L, H).

Many OFC neurons (463 out of 649 neurons, p<0.05) were responsive to the learned visual cues. The activity of many neurons was related to a single variable. Figure 4 shows the cue-period activity of some example neurons. The neuron illustrated in Figure 4A had a higher firing rate when the monkey expected to receive 3 drops of juice relative to 1 drop of juice. Its activity did not differ between volatile and stable cues, or between certain and uncertain cues (Figure 4A). Thus, this neuron’s activity was modulated only by the magnitude of the reward predicted by the monkey. The second example neuron showed greater activity to the cues that signaled volatile reward outcomes (F, S, PF, PS) relative to those that signaled nonvolatile outcomes (H, L, PH, PL). Interestingly, its activity did not appear to be modulated by reward magnitude or certainty (Figure 4B). The third example neuron showed a higher firing rate for the cues that signaled uncertain outcomes (PH, PL, PF, PS) relative to those that signaled certain outcomes (H, L, F, S), but its activity was not modulated by reward magnitude or volatility (Figure 4C). Finally, we also found neurons such as the one shown in Figure 4D that had a mixed tuning.

These examples suggest there was a diversity of neurons in the OFC that reflected the different aspects of reward that we studied. We used a linear regression model (see Materials and Methods) to quantify the neurons’ tuning during the cue period. Out of the total number of 649 neurons that we recorded, we found 158 neurons (24.3%) were tuned to reward magnitude, 167 neurons (25.7%) were tuned to volatility, and 154 neurons (23.7%) were tuned to certainty (Figure 5D). Only a small proportion of neurons—although more than predicted by chance on the assumption that the tuning of the three aspects was independent—was tuned to more than one reward aspect (reward magnitude and volatility: 64 neurons (9.9%); reward magnitude and certainty: 48 neurons (7.4%); volatility and certainty: 71 neurons (10.9%); all three: 29 neurons (4.5%); all ps<0.05, permutation test). Notably, a relatively large proportion of neurons (238/649) was tuned only to certainty or volatility, but not to reward magnitude.

In sum, we identified three groups of OFC neurons active during the cue period, one each that selectively signaled reward magnitude, certainty and volatility (Table 1). The latency of the population encoding of each tuning property was similar (155 ms for certainty and volatility, and 160 ms for reward magnitude), suggesting the encoding of these three attributes was simultaneous (Figure 5A). In addition, we calculated all recorded OFC neurons’ tuning indices for each attribute. The reward magnitude, volatility or certainty indices were not correlated with one another (Figure 5B and C). These results indicate that the encoding of these three properties was orthogonal in the OFC.

The neuronal responses to cue presentation were transient. We plotted the population responses related to reward magnitude during the cue presentation period, the period immediately after cue offset, and the period before button release signaled the monkey’s estimation of reward magnitude. For neurons that encode reward magnitude, the encoding of reward magnitude could be seen as early as 160 ms after cue onset, and it disappeared shortly after cue offset (Figure 6A). Although the monkey had to remember the predicted reward magnitude during the delay period to perform the task, these OFC neurons did not seem to be involved in maintaining the information.

In many studies, OFC neurons have been found to carry signals about expected reward magnitude from the cue period to the time of reward delivery. It was therefore surprising that our OFC neurons fell quiet after the cue offset. Accordingly, we also evaluated the encoding of reward magnitude later in the trial, near the time of reward delivery. During the 500-ms period leading to the release of the button that indicated the monkeys’ responses (1 or 3 second hold), 244 neurons (37.6%) encoded the predicted reward magnitude (Table 1, Figure 6B). This is a significant increase from the 158 neurons (24.3%) that encoded reward magnitude during the cue period. As can be deduced from the lack of encoding of reward magnitude during the delay period, the pre-release activity of these neurons was not a continuation of their activity during the early cue period. As a group, these neurons only exhibited small cue responses, and the responses were not modulated by reward magnitude during the cue period (Figure 6B). In fact, only 68 neurons (10.5%) encoded reward magnitude during both the cue and the pre-release period. The number of neurons that encoded predicted reward magnitude in both periods is not significantly different from that expected by chance (p<0.05, permutation test). In addition, 25 of these 68 neurons actually reversed their signs, e.g., showed greater activity for greater reward magnitude in the cue period but less activity for longer holding periods in the pre-release period. Thus, we concluded that the reward magnitude encoding during the pre-release period was represented separately in the OFC from the encoding in the early cue period.

## Discussion

Stimulus-reward association is often uncertain and volatile in the real world. The same sensory event may lead to different outcomes at different times. Conversely, the same outcome may be interpreted differently. Previous studies have focused on how probabilities can be incorporated into the estimation of reward or value, for instance, by multiplying probability and magnitude. Knowledge of the state or condition of outcomes being certain or volatile is important in its own right, however, and needs to be taken into account for tracking reward information successfully in a complex environment. Here we demonstrate that neurons in the OFC encode whether outcomes are certain and/or volatile in addition to encoding reward magnitude. More importantly, they do so in an unexpected manner.

First, we found that the certainty and volatility aspects of stimulus-reward associations were represented separately from the magnitude of anticipated reward in the OFC. One group of neurons indicated whether the reward associated with a visual stimulus was certain or not, another group indicated whether it was volatile, and a third group encoded how much reward could be received. The majority of neurons in the certainty and volatility groups were not sensitive to reward magnitude. Even though there were neurons with activity profiles that reflected mixed tuning, they were a minority. These results are in contrast with previous findings that many OFC neurons encoded multiple variables such as reward and cost during decision making (Kennerley and Wallis 2009). One possible reason for this apparent discrepancy is that, in our study, certainty and volatility have to be considered separately for the monkey to adapt to the changing environment; therefore, our task design might have promoted independent coding of these factors. Another difference between the studies relates to whether the neurons encoded reward probability *per se* (Kennerley and Wallis 2009), or instead reflected the categorical knowledge that a given cue varied in the dimension of certainty or volatility (present study).

Second, there are roughly equal numbers of neurons that encode certainty, volatility, and magnitude. As reward magnitude is the more basic property of stimulus-reward associations, and certainty and volatility can be inferred only from reward magnitude over history, it is surprising that the three are represented at similar strength in the OFC. This may indicate that the OFC, instead of calculating reward value, *per se*, is a place where different aspects of reward information converge, allowing it to establish stimulus-reward associations and to optimize choice behavior.

Finally, we note that the representation of reward information during the visual stimulus presentation was transient. The activity of the neurons that were tuned to reward magnitude, certainty and volatility dropped back to the baseline quickly after the cue offset. They did not show a sustained activity that might support decision-making at later times. In this aspect, these OFC neurons resemble neurons in early sensory areas. As indicated earlier, this is unlike several other studies that have reported sustained activity from the cue period through to receipt of reward, including studies in which the cue was presented briefly, as done here (Wallis and Miller 2003; Raghuraman and Padoa-Schioppa 2014). Perhaps the transient activity reflects the fact that the monkeys made their prediction of the reward magnitude, which could be considered a ‘choice’, at the termination of the cue period. If so, there was no need to retain the information about reward magnitude, as the response had been selected.

Another group of OFC neurons was active during the period immediately before monkeys indicated their estimation of reward size for that trial. In contrast to the neurons that were active during the cue period, they mostly encoded the expected reward magnitude. A small number of neurons were active in both task epochs (cue and pre-release periods). However, even these neurons did not show sustained activity in the period between, and their reward magnitude encodings in the two periods were uncorrelated. Therefore, their responses during the pre-release period are likely due to recurrent projections. We do not know if the latter group of OFC neurons received information directly from those active during the cue period, from another group of neurons within the OFC, or from neurons in a different brain region.

Notably, our results are consistent with the proposal that there is a representation of task space in the OFC (Wilson et al. 2014). The concept of task space is key to the reinforcement learning framework. Appropriate learning from feedback requires knowledge of the task structure. In the present experiment, certainty and volatility information associated with each visual stimulus can be considered part of the task structure that is necessary for guiding optimal choice behavior. Remarkably, we found not only that there are representations of the three properties of reward we studied in the OFC, but also that the proportions of neurons encoding those properties were similar, reflecting the fully balanced design of the task. That is, magnitude, certainty, and volatility were equally represented by the stimuli used in the experiment. It is reasonable to speculate that our findings were driven by this design, as one might expect if OFC neurons were encoding task space. The finding of uncorrelated encoding of reward magnitude in the cue and pre-release periods also is consistent with this idea. During the cue period, all aspects of reward are important state parameters, of which reward magnitude is only one. During the pre-release period, the most important state is the reward magnitude, which is useful for the monkey to generate its motor response and to evaluate feedback regarding the accuracy of estimated reward size. This would seem to account for why encoding of reward magnitude dominates the pre-release period.

Given that response hold time (short or long) is confounded with reward magnitude, an alternative account is that the neurons active in the cue period are encoding motor responses, either in addition to reward magnitude or instead of it. This seems unlikely, however, as previous studies suggest there is a paucity of action related signals in the OFC neural activity (Wallis and Miller 2003; Cai and Padoa-Schioppa 2014). Furthermore, the selectivity during the cue period died out after the cues were turned off, suggesting that the cue period activity is probably not related to motor responses.

We are yet to understand how knowledge of certainty and volatility is integrated with reward magnitude. Because OFC neurons separately encoded reward variables, our results suggest that integration occurs in a brain structure other than the OFC. A likely candidate is the dorsolateral prefrontal cortex, which has been linked to many cognitive functions, including especially those that require multiplexing (Duncan 2010). Another candidate region for integration is the anterior cingulate cortex (ACC), which has been suggested to play an important role in detecting errors, monitoring conflicts, and generating reward predictions in general (Botvinick et al. 2004; Hayden and Platt 2010; Kolling et al. 2016; Shenhav et al. 2016). In our task, monkeys had to monitor rewards to guide adaptive response selection (long or short hold). Signals guiding decisions to switch from a default response have been found in dorsal ACC (Hayden et al. 2011; Boorman et al. 2013) and may well be relevant for task performance in the present study. Other brain areas, including the supplementary eye field, have also been shown to encode certainty (So and Stuphorn 2012). Lastly, the OFC may interact with the basal ganglia to use knowledge of certainty and volatility to optimize choices. The two structures together form an OFC-basal ganglia loop that plays an important role in processing reward information (Haber and Behrens 2014).

Although the three variables we considered all contribute to the calculation of reward value, the current study did not address how reward value is calculated. Such a calculation requires additional information, such as temporal delay and effort. In the present study, the exact value of the outcome is not important. The monkeys were not making choices based on reward value. Instead, they were required to report the current state of the reward cues based on knowledge of their certainty and volatility. How certainty and volatility affect monkeys’ value-based decision making invites empirical investigation.

Finally, our study adopted a categorical approach to study certainty and volatility. Different levels of certainty and volatility are not captured by this approach. To properly study the cue-reward associations, we interleaved all cue types in the task. The number of conditions has to be small enough to record sufficient trials in each condition for each neuron. Future studies may focus on one aspect of the reward, either certainty or volatility, to study how different levels of these features are represented in the brain.

In summary, we show that OFC neurons encode categorical knowledge of reward certainty and volatility associated with visual stimuli. This not only adds to the rich literature that demonstrates the importance of the OFC in learning stimulus-reward associations, but also shows that such associations extend to more abstract features of reward. It is an important step towards a full understanding of the cortical mechanisms of reward processing and reward-based decision making.

## Acknowledgments

We thank Peter Rudebeck and Andrew Mitz for their help in all phases of the study, and Ping-yu Chen for histological support. The authors declare no competing financial or nonfinancial interests.

